# Highly pathogenic avian influenza A (H5N1) in marine mammals and seabirds in Peru

**DOI:** 10.1101/2023.03.03.531008

**Authors:** Mariana Leguia, Alejandra Garcia-Glaessner, Breno Muñoz-Saavedra, Diana Juarez, Patricia Barrera, Carlos Calvo-Mac, Javier Jara, Walter Silva, Karl Ploog, Lady Amaro, Paulo Colchao-Claux, Marcela M. Uhart, Martha I. Nelson, Jesus Lescano

## Abstract

Highly pathogenic avian influenza (HPAI) A/H5N1 viruses (lineage 2.3.4.4b) are rapidly invading the Americas, threatening wildlife, poultry, and potentially evolving into the next global pandemic. In November 2022, HPAI arrived in Peru, where massive pelican and sea lion die-offs are still underway. We report complete genomic characterization of HPAI/H5N1 viruses in five species of marine mammals and seabirds (dolphins, sea lions, sanderlings, pelicans and cormorants) sampled since November 2022. All Peruvian viruses belong to the HPAI A/H5N1 lineage 2.3.4.4b, but they are 4:4 reassortants where 4 genomic segments (PA, HA, NA and MP) position within the Eurasian lineage that initially entered North America from Eurasia, while the other 4 genomic segments (PB2, PB1, NP and NS) position within the American lineage (clade C) that was already circulating in North America. These viruses are rapidly accruing mutations as they spread south. Peruvian viruses do not contain PB2 E627K or D701N mutations linked to mammalian host adaptation and enhanced transmission, but at least 8 novel polymorphic sites warrant further examination. This is the first report of HPAI A/H5N1 in marine birds and mammals from South America, highlighting an urgent need for active local surveillance to manage outbreaks and limit spillover into humans.

## INTRODUCTION

The recent emergence of highly pathogenic avian influenza (HPAI) H5N1 viruses in mammals and birds in the Americas^1,2^ presents a severe threat to wild and endangered species, to poultry production^3,4^, and to public health when the virus spills over into humans^3,5^. The H5N1 (clade 2.3.4.4b) virus arrived in North America in late 2021 from Eurasia and spread across the continent in wild birds, spilling over into poultry farms and infecting an alarming number of wild terrestrial mammals, including fox, skunk, bear, bobcat, and raccoon. In October 2022, an apparent mink-to-mink transmission of clade 2.3.4.4b H5N1 viruses in Spain^6^ further heightened concern that the avian virus was adapting to mammals and that an H5N1 global pandemic in humans could be approaching.

In November 2022, Peruvian pelicans (*Pelecanus thagus*) along coast and in offshore islands began experiencing a mass die-off^7,8^. HPAI A/H5N1 was identified in samples collected from dead birds, resulting in the country declaring a sanitary alert^7^. This was followed by several spillover events into other domestic and wild birds, including zoo animals and wild raptors^4,8^. By the beginning of 2023, the Peru outbreak had spread to marine mammals, particularly affecting the South American sea lion (*Otaria flavescens*), which also began to experience a mass die-off^9^.

The Pacific coast of Peru hosts a rich biodiversity of marine mammals and seabirds^10^. The ecosystem is home to large populations of South American sea lion, “guano birds’’ like the Guanay cormorant (*Phalacrocorax bougainvillii*), the Peruvian booby (*Sula variegata*), and the Peruvian pelican, and endangered birds like the Humboldt penguin (*Spheniscus humboldti*). The region also serves as stopover points and feeding grounds for diverse avian migratory species, including the Franklin’s gull (*Leucophaeus pipixcan*) and several species of sandpipers (*Calidris* spp.)^11,12^ that travel south from the northern hemisphere during the boreal winter. Here we report detection, genomic characterization, phylogenetic analysis, and mutation analysis of HPAI A/H5N1 viruses identified in marine mammals (sea lion and common dolphin) and seabirds (pelican, cormorant and sanderling) sampled along the coast of Peru since November 2022.

## MATERIALS AND METHODS

### Sample collection and pre-processing for influenza A

Starting on November 22, 2022, samples were collected opportunistically from both live and dead animals by trained veterinarians from the Peruvian wildlife service (Servicio Nacional Forestal y de Fauna Silvestre, SERFOR) and the NGO Wildlife Conservation Society Perú, under permit XX, using full personal protective equipment and following standard protocols for cleaning, disinfection, and disposal of hazardous waste^13^. Some severely clinically ill animals were humanely euthanized^14^. Briefly, animals were chemically immobilized by intramuscular administration of ketamine and xylazine, followed by intravenous or intracardiac injection of T-61™ (MSD Animal Health USA)^14^. Samples and tissues were collected into cryovials containing 0.5 mL of DNA/RNA Shield (Zymo R1200 125) or viral transport media (VTM), and transported at 4°C within 1-10 days of collection to the laboratory for testing. Nucleic acids were extracted using Quick-DNA/RNA Viral Extraction Kits (Zymo D7021) and tested for influenza A by RT-qPCR using CDC protocols^15^ on a Bio-Rad CFX96 instrument.

### Influenza A subtyping

Samples positive for influenza A by RT-qPCR were subtyped using a combination of directed amplification with universal primers targeting conserved genomic regions^16^, followed by next-generation sequencing (NGS). Briefly, RNA samples were reverse transcribed using Superscript IV (Invitrogen 18090050) and amplified using Q5 High-fidelity DNA polymerase (NEB M0491L). Amplification products were prepared into barcoded sequencing libraries using DNA Prep Kits (Illumina 20060059) and Nextera DNA UD Indexes (Illumina 20027215). The resulting libraries were quality controlled using High Sensitivity DNA kits (Agilent 5067-4626) on a Bioanalyzer 2100 instrument.

Libraries were normalized to 4nM each, pooled, re-quantified using Qubit 1x dsDNA HS Kits (Invitrogen Q33230), and sequenced using High Output Sequencing Kits (Illumina FC-420-1003) on an Illumina MiniSeq instrument.

### Bioinformatics processing

Illumina paired-end raw sequence data was pre-processed to trim sequencing adaptors and filter out low quality/low complexity reads (Phred scores <Q20, 75 bp minimum length) using Geneious Prime 2023.0.4 and BBDuk^17^. Pre-processed reads were then filtered by reference-mapping to various HA (H1, H2, H3, H5, H7) and NA (N1, N2, N3, N5, N7) references (Accession Numbers: NC_026433.1, NC_007374.1, NC_007366.1, NC_007362.1, NC_026425.1, AF144304.1, NC_007382.1, OP806485, MF046172.2, OP723829.1), and to single references in the case of all other segments (Accession Numbers MT624412.1, MN254461.1, KY635563.1, MT825070.1, MW110227.1, MT982385.1). Filtered reads were then re-assembled *de-novo* using SPAdes^18^ to generate complete genomes whenever possible, and to further confirm subtyping by BLAST. All sequences have been deposited in GenBank (Accession Numbers XX).

### Phylogenetic and mutation analysis

HA and NA reference sequences were obtained from GenBank and GISAID, and together with the sequences generated here, they were concatenated and aligned using MAFFT^19^. Maximum likelihood (ML) trees were prepared with FastTree^20^ incorporating a general time-reversible (GTR) model of nucleotide substitution with a gamma-distributed rate variation among sites. Trees were annotated using the WHO/OIE/FAO nomenclature for highly pathogenic avian influenza A H5^21^. To place the Peruvian viruses in a global context, we downloaded (on February 14, 2023) an additional background dataset of influenza A virus genomes that included all sequences from avian and mammalian H5 viruses submitted to GISAID since January 1, 2021. Partial sequences were excluded.

Phylogenetic relationships were inferred for each of the eight genome segments using the ML methods available in IQ-Tree 2^22^ with a GTR model and a GAMMA distribution as described above. Due to the size of the dataset, we used the high-performance computational capabilities of the Biowulf Linux cluster at the National Institutes of Health (http://biowulf.nih.gov). To assess the robustness of each node, a bootstrap resampling process was performed with 1000 replicates. Finally, mutation analysis was done using the CDC H5N1 genetic changes inventory for SNP analysis and various other previously published mutations of concern^23,24^.

## RESULTS

### Detection of HPAI positive samples in mammals and seabirds in Peru

Starting in November 2022, we collected swabs of external orifices and internal organ tissues from six species of marine mammals and seabirds (common dolphin (*Delphinus delphis)*, South American sea lion *(Otaria flavescens)*, sanderling *(Calidris alba)*, Peruvian pelican *(Pelecanus thagus)*, Guanay cormorant *(Phalacrocorax bougainvillii)* and Humboldt penguin *(Spheniscus humboldti)*) in regions representing the northern (Piura), central (Lima) and southern (Arequipa) coast of Peru (Supplemental Table 1, Figure1). All animals sampled were either deceased or manifested clear signs of disease, including respiratory, digestive, and/or neurological symptoms indicative of acute encephalitis. Seabirds exhibited disorientation, ataxia, circling, nystagmus, torticollis, congested conjunctivae and dyspnea; sea lions exhibited disorientation, ataxia, circling, “stargazing” posture, copious nasal discharge, sialorrhea, dyspnea and seizures; the common dolphin was found recently deceased.

**Figure 1:**
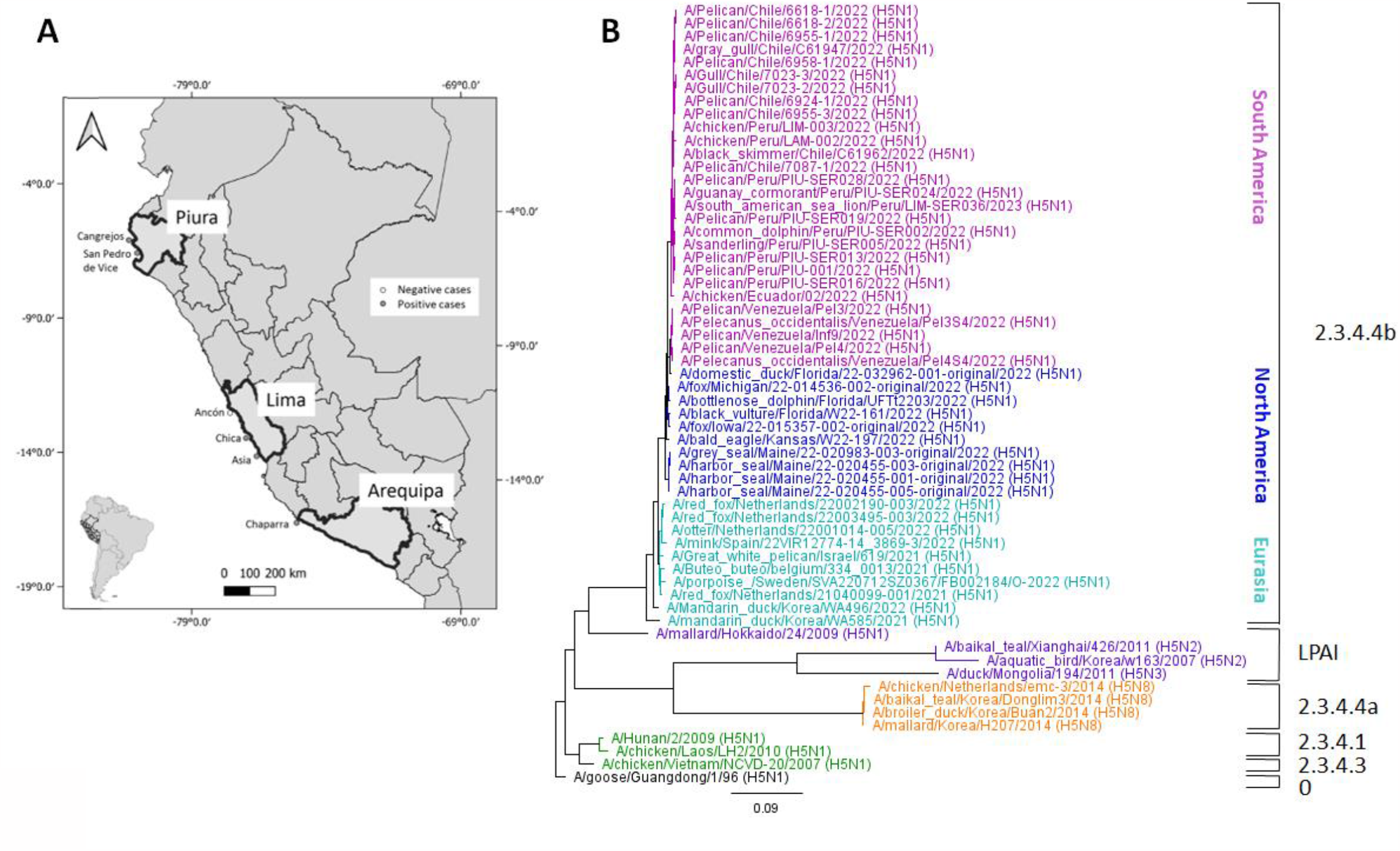
Collection sites and basic phylogeny of HPAI A/H5N1. (A) Map of Peru showing collection sites representative of the northern, central and southern regions. Mass die-offs of both birds and mammals are ongoing along the entire coast of Peru. (B) Phylogenetic tree for concatenated HA and NA segments inferred using ML methods in FastTree2 (GTR+GAMMA) and rooted to ancestral A/goose/Guangdong /1/1996. Clade designation is based on WHO/OIE/FAO H5 Evolution Working Group nomenclature. Clade O: Ancestral Root. LPAI: Low pathogenic Avian Influenza strains.

To date, we have tested a total of 63 samples from 24 individuals by RT-qPCR, confirming 10 individuals (1 dolphin, 3 sea lions and 6 seabirds), plus 1 pool of 5 sea lions, as positive for influenza A. Positives were subjected to NGS for subtyping and to generate full genomes for phylogenetic and mutational analysis. We confirmed 8 individuals as positive for HPAI A/H5N1 (1/1 dolphin, 1/3 sea lions and 6/6 seabirds). It was not possible to type the remaining positives, as they exhibited very low viral loads (Supplemental Table 1) or extensive signs of nucleic acid degradation, likely associated with decomposing tissues in deceased animals (not shown). Of the 8 individuals that yielded quality sequence data, we generated complete sequences for most genomic regions in most samples (Table 1 and Supplemental Table 1). All sequences have been deposited in GenBank (Accession Numbers XX).

**Table 1:** Eight Peruvian HPAI A/H5N1 genomes sequenced in this study.

| Virus name | Species | Sample | Found | Sex | Age | Date | Location | Accession |
| --- | --- | --- | --- | --- | --- | --- | --- | --- |
| A/common dolphin/Peru/PIU-SER002/2022 | *D. delphis* | Rectal | Dead | M | Adult | 2022-11-22 | Piura (north) |  |
| A/sanderling/Peru/PIU-SER005/2022 | *C. alba* | Trachea | Euthanasia | n/a | Juvenile | 2022-11-22 | Piura (north) |  |
| A/pelican/Peru/PIU-SER013/2022 | *P. thagus* | Brain | Dead | M | Adult | 2022-11-23 | Piura (north) |  |
| A/guanay cormorant/Peru/PIU-SER024/2022 | *P. bougainvillii* | Brain | Euthanasia | F | Adult | 2022-11-24 | Piura (north) |  |
| A/pelican/Peru/PIU-SER019/2022 | *P. thagus* | Brain | Dead | M | Adult | 2022-11-24 | Piura (north) |  |
| A/pelican/Peru/PIU-SER028/2022 | *P. thagus* | Brain | Euthanasia | F | Adult | 2022-11-24 | Piura (north) |  |
| A/pelican/Peru/PIU-SER016/2022 | *P. thagus* | Lung | Euthanasia | F | Adult | 2022-11-24 | Piura (north) |  |
| A/South American sea lion/Peru/LIM-SER036/2023 | *O. flavescens* | Lung | Abortion | M | Abortion | 2023-01-23 | Lima (central) |  |

### Eurasian-American lineage reassortants

To further characterize our Peruvian HPAI A/H5N1 isolates we carried out phylogenetic analyses based on concatenated sequence data of the HA and NA genomic regions. The analysis included all HPAI A/H5N1 reference sequences for South America available to date in public repositories, plus other references representative of the H5N1 lineage going back to the A/H5N1 goose/Guangdong strain originally identified in 1996^25^. Our analysis confirms that all samples sequenced are closely related and cluster within the HPAI A/H5N1 lineage 2.3.4.4b (Figure 1). Additional phylogenetic analyses conducted separately for each genome segment revealed that our eight Peruvian HPAI A/H5N1 isolates are reassortants (Figure 2, Table 2). The Peruvian viruses are positioned in the Eurasian lineage in trees inferred for the PA, HA, NA and MP segments (Pattern A), clustering with the North American viruses that were first introduced from Eurasia in late 2021. In contrast, the Peru viruses are positioned in the American lineage for trees inferred for the PB2, PB1, NP and NS segments (Pattern B), which is evidence of a 4:4 reassortment event (Figure 2 for representative PB2 and PA trees, and Supplementary Figures for the remaining six segment trees). Most of the H5N1 viruses observed since the summer of 2022 in the Americas are reassortants with various combinations of segments from the Eurasian and American lineages (Figure 3A). The Eurasian lineage H5N1 virus that originally invaded North America in 2021 has reassorted multiple times with the endemic American lineage since arriving in North America, as evidenced by multiple reassortant clades positioned within the American lineage (Figure 2: A-E). The Peru viruses are positioned in clade C on the PB2, PB1, NP and NS trees and their 4:4 reassortment pattern is referred to as pattern “R6” (Table 2).

**Table 2:**
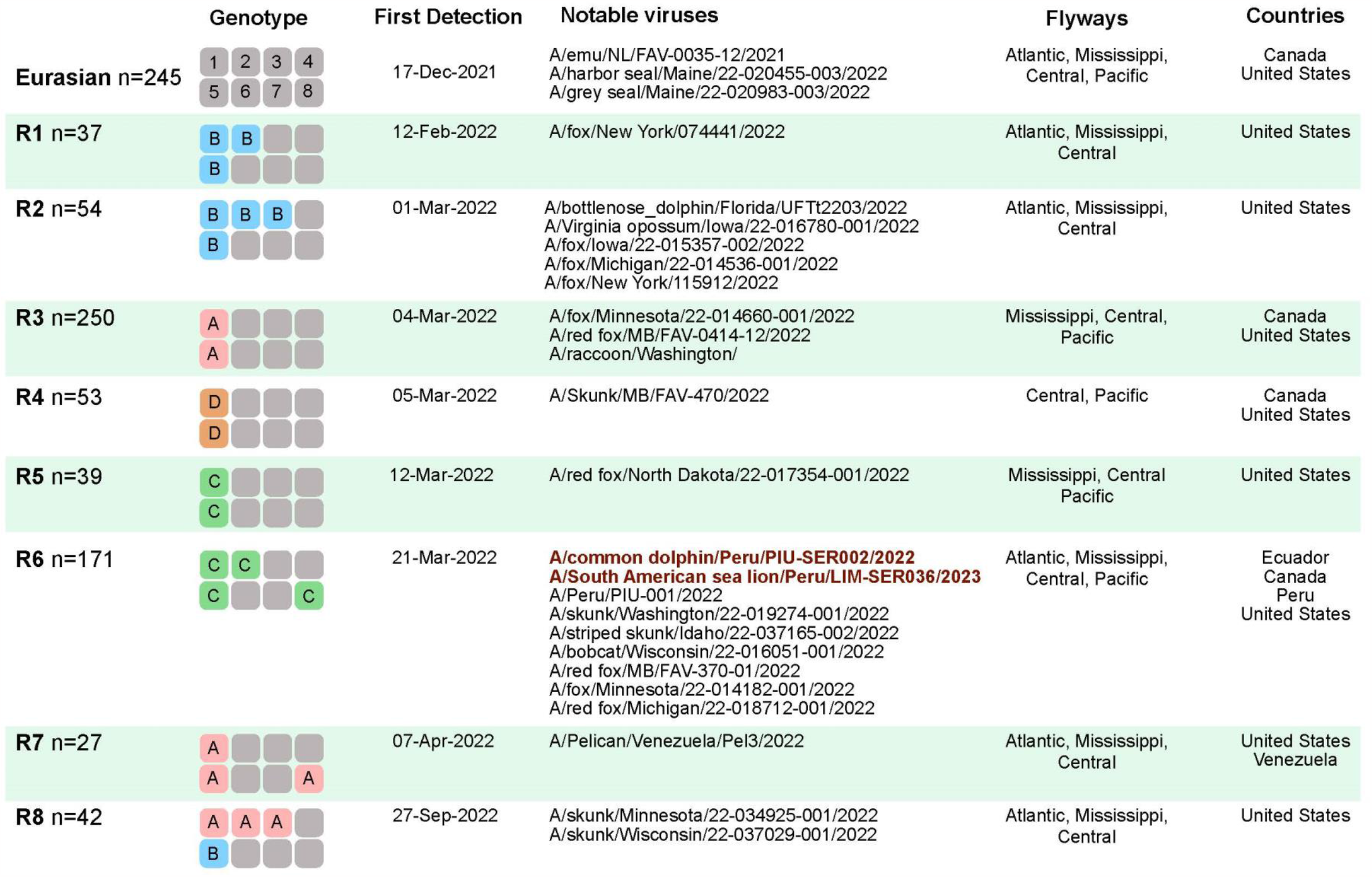
H5N1 genotypes identified in the Americas, 2021-2023. Each genotype box represents one of eight genome segments (1-8), shaded by lineage: grey = Eurasian lineage; pink = American lineage (clade A); blue = American lineage (clade B); green = American lineage (clade C); orange = American lineage (clade D).

**Figure 2:**
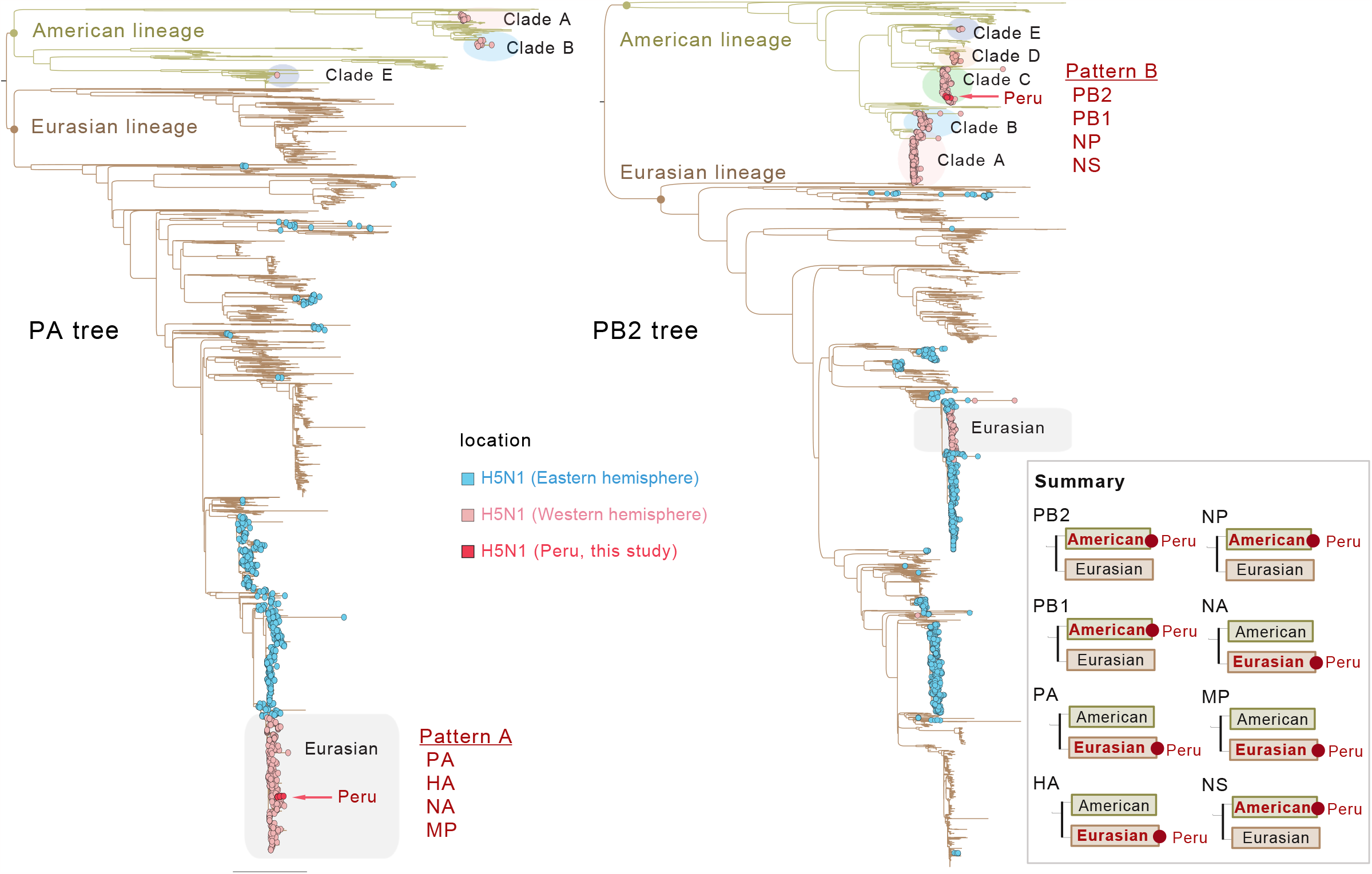
Global H5N1 phylogeny for the PA (Pattern A) and PB2 (Pattern B) segments. Phylogenetic trees for AIVs collected globally (PA n=5,438; PB2 n=5,464) and submitted to GISAID since January 1, 2021, inferred using ML methods. Branches shaded by AIV lineage; tips shaded by AIV subtype and location. H5N1 viruses from the 2021-2023 Western hemisphere outbreak are shaded pink and labeled with a grey box (original introduction of Eurasian-origin H5N1) or colored circles (clades A-E, American lineage reassortants). The 8 Peruvian viruses described here are shaded red and indicated with arrows. These viruses are 4:4 reassortants, where 4 segments (PA, HA, NA, MP) position in the Eurasian lineage (Pattern A, using the PA tree), and 4 segments (PB2, PB1, NP, NS) position in Clade C of the American lineage (Pattern B, using the PB2 tree).Trees for all eight segments, including bootstrap values and scale bars, are provided in the Supplementary Materials. A summary of all tree patterns is provided in the lower right box.

**Figure 3:**
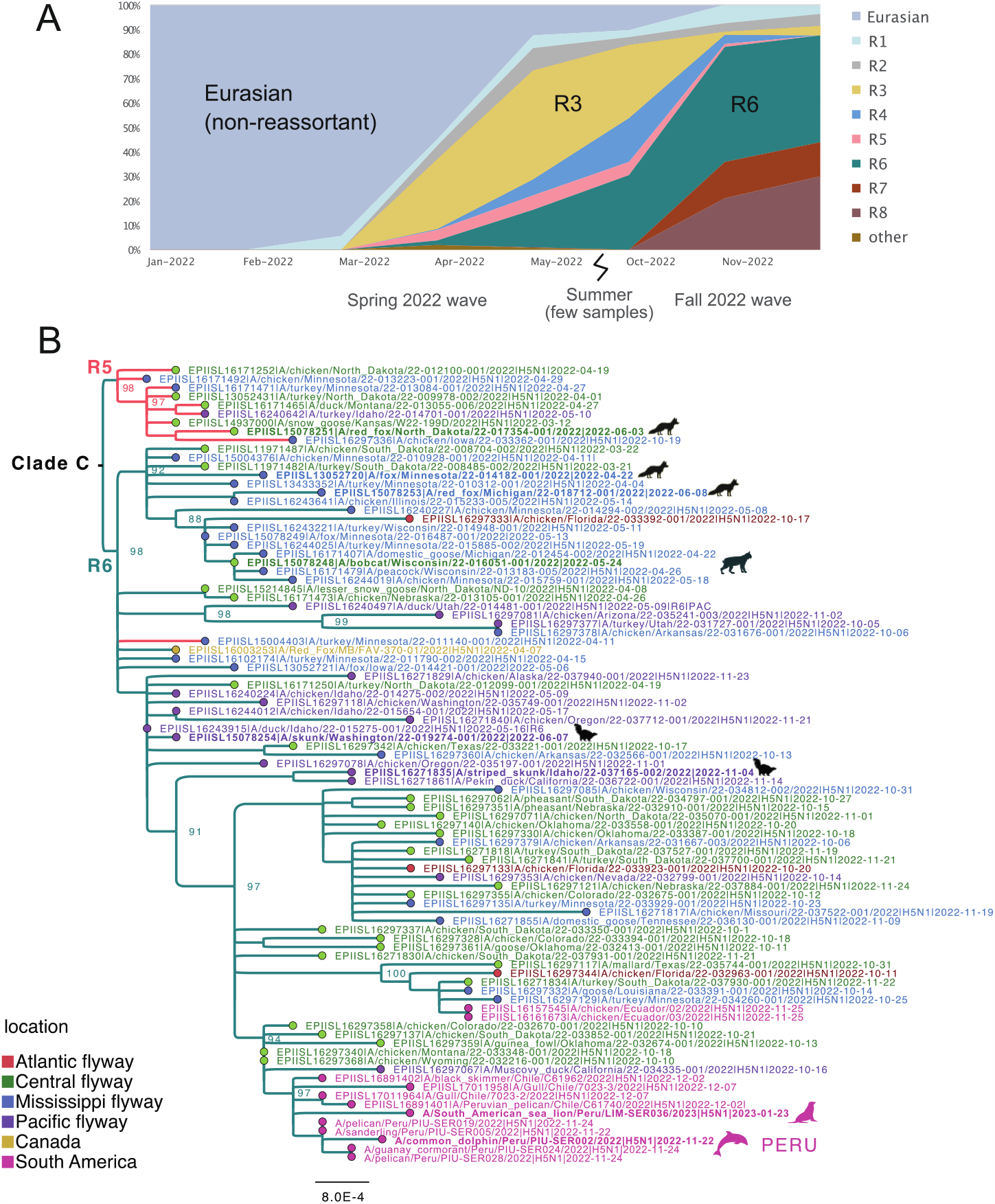
Spread of reassortant “R6” viruses. (A) Proportion of H5N1 viruses sequenced in the Americas between 2021-2023 that belong to different genotypes (Table 2) over time. (B) Phylogenetic tree inferred using ML methods for 87 representative H5N1 viruses from clade C (see Figure 2). Branch lengths are drawn to scale and shaded by reassortant genotype (R5 = pink; R6 = teal, Table 2). Tips are shaded by location/flyway. Bootstrap values provided for key nodes. Viruses collected from mammals are indicated by cartoons.

R6 reassortants were first detected in North American poultry in March 2021 and became dominant among sequenced viruses in the Americas during the autumn of 2022 (Figure 3A). The population genetics of H5N1 in the Americas underwent a shift during 2022, as reassortants displaced the original non-reassortant Eurasian H5N1 virus, which has not been detected in the Western hemisphere since June 2022. All currently detected H5N1 reassortants in the Western hemisphere still retain Eurasian HA, NA, and MP segments, but their PB2 and NP segments belong to the American lineage. Eurasian and American PB1, PA, and NS segments continue to co-circulate in the Americas in different reassortant backgrounds. Viruses isolated from poultry in Ecuador in November 2022 (e.g., A/chicken/Ecuador/02/2022) have the same R6 reassortant genotype as our Peru viruses, but are positioned in a different section of clade C (Figure 3B), meaning the South American H5N1 viruses are not monophyletic and represent independent viral introductions from birds migrating to South America from the north. The viruses isolated from pelicans in Venezuela in November 2022 have a different reassortant genotype (R7) that is primarily found in the eastern US, including in Florida, and represents a third independent viral introduction from North to South America (Table 2). Four H5N1 viruses from Chile (e.g., A/gull/Chile/7023-3/2022) cluster with our Peru viruses and have the same R6 genotype, evidence of spread of the Peruvian outbreak into Chile. This clade of Peru and Chile viruses appears to descend from a single viral introduction from North America, which then transmitted in Peru between avian and mammalian species (pelican, cormorant, sandpiper, dolphin and sea lion), and then onward to Chile. At this point however, the direction of transmission among these different avian and mammalian species within Peru is not possible to infer with the available data.

### Mutation analysis

We also performed a detailed SNP and mutational analysis to identify amino acid changes potentially linked to increased virulence, transmission, or mammalian host adaptation, and to assess if we could identify specific differences between host species (mammals vs. birds). We identified more than 70 variable sites spread across all genomic regions (Figure 4, Supplementary Table 2), including 40 that have been previously reported as linked to specific phenotypes, and 30 that remain uncharacterized. Of the 40 previously reported polymorphic sites, 8 were in PB2 (L89V, G309D, T339K, Q391E, R477G, I495V, K627E, A676), 1 in PB1 (N375S), 1 in PB1-F2 (N66S), 4 in PA (I61M, T85A, A515T, T608S), 12 in HA (E75K, S123P, S133A, S154N, S155D, T156A, K189N, K218Q, S223R, L269V,Q322L, R325K), 1 in NP (S450N), 5 in M1 (N30D, I43M, N85S, K101R, T215A) and 7 in NS1 (P42S, D92E, L98F, I101M, D134N, V144A, A218E). These changes have been linked to a number of relevant phenotypes, including altered polymerase activity and replication efficiency (usually enhanced), increased virus binding to α2-3 and α2-6, enhanced transmission, and increased virulence and pathogenicity, including in mammals^23,24^. We also identified an additional 30 sites spread across all genomic segments (Figure 4, Supplementary Table 2), including 4 in PB2 (I463V, L464M, V478I, I616V), 5 in PB1 (T59S, E264D, L378M, G399D, K429R), 11 in PBP1-F2 (T7I, S12L, N17S, R21K, Y42C, P48Q, Q54R, I55T, Y57S, W58L, G70E), 1 in PA (M441V), 2 in HA (I511V, V533M), 1 in NP (F230L), 2 in NA (L269M, S339P), 2 in M1 (N87T, A200V), 1 in M2 (R61G) and 2 in NS1 (C111S, G166D). We did not find mutations in PB2 (E627K, D701N, K702R) that have been specifically linked to mammalian host adaptation and enhanced transmission^24,26^. We also did not find mutations specific to mammalian or bird strains that might support mammalian host adaptation, but we did find a small number of isolated mutations in single birds (not shown) or mammals (Supplement table 2). The virus from sea lion contains 7 unique mutations that may warrant further observation: 2 in PB2 (T215M and N715T), 1 in PA (M86I), 1 in HA (A496S), and 3 in NP (M222L, A428T and R452K). Given that these were not present in the other mammal sequenced here, sampling of additional individuals will be needed to assess their significance. The remaining 30 uncharacterized variable sites are present across all species sampled, and in most cases, they are restricted only to sequences reported in Latin America, and in a few cases to select sequences from North America (Supplementary Table 2), which indicates that the virus is indeed changing as it travels south from the northern hemisphere.

**Figure 4:**
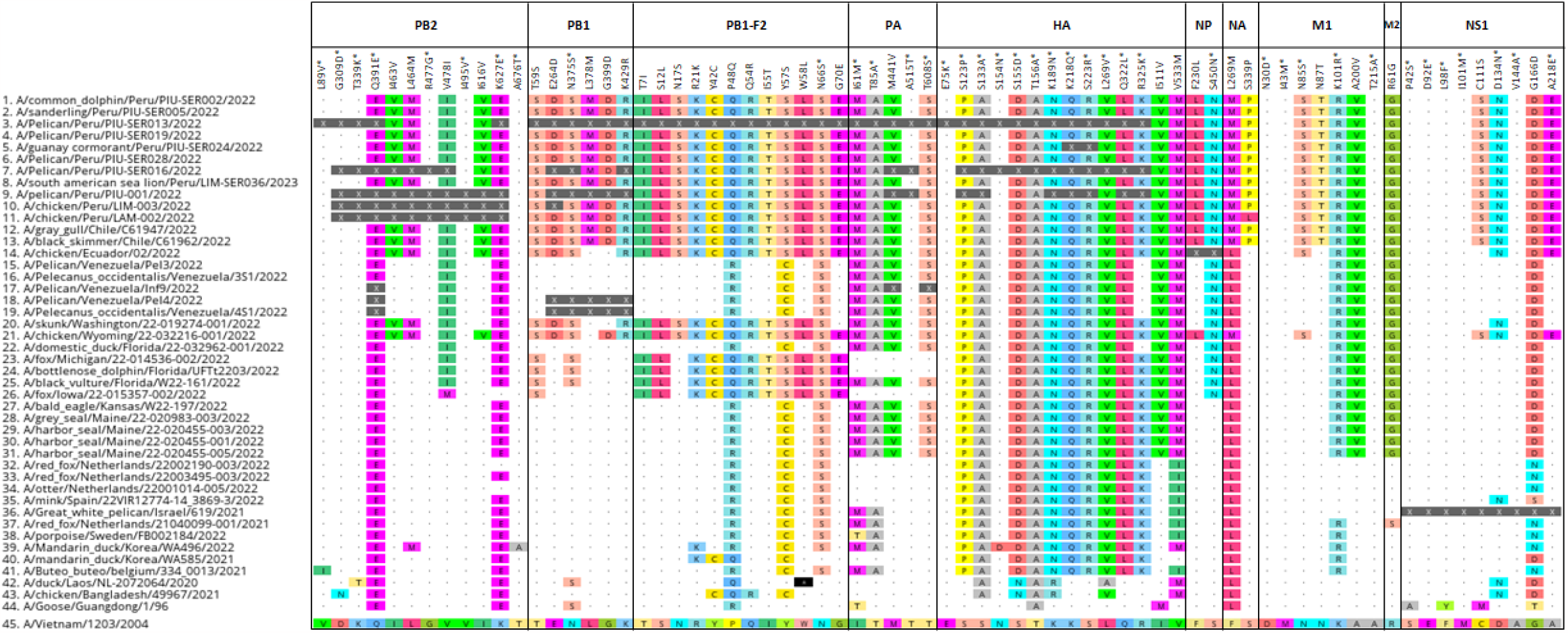
SNP and mutational analysis of Peruvian HPAI a/H5N1 viruses. Additional reference sequences used are available through GenBank and GISAID. A total of 70 variable sites were identified relative to the original A/H5N1 goose/Guangdong reference from 1996 and the A/Vietnam/1203/2004 reference used to annotate amino acid positions in the CDC inventory^23^. 40/70 sites identified have been previously reported as associated with mutations of interest and are shown with an asterisk (*). The remaining 30/70 sites have not been previously characterized. Dots represent amino acids identical to those present in annotation reference A/Vietnam/1203/2004.

## DISCUSSION

For decades, South America’s relative geographic isolation has largely insulated its fragile coastal ecosystems and poultry industry from the HPAI outbreaks that periodically ravage US and Mexican farms to the north. But the newest variant of HPAI A/H5N1, clade 2.3.4.4b, is spreading faster, causing mass mortality in wildlife, infecting wild mammals, and invading countries like Peru that had remained HPAI-free for decades. The arrival of HPAI in regions with less experience managing highly pathogenic viruses in wildlife and poultry is concerning. Here, we rapidly established new surveillance partnerships between government and academia to respond to the mass die-offs of Peruvian pelicans and South American sea lions currently underway in Peru. We confirmed the presence of HPAI A/H5N1 clade 2.3.4.4b in both pelicans and sea lions, as well as in Guanay cormorans, sanderlings and dolphins, and further surmised that HPAI A/H5N1 is the likely causative agent of the mass wildlife die-offs currently underway in Peru. We suspect that direct HPAI transmission between sea lions could be occurring, rather than independent spillovers into sea lions from avian sources, but additional sequence data and analysis will be required to assess this. We report 40 variable sites previously linked to altered polymerase activity and replication efficiency, increased virus binding to α2-3 and α2-6, enhanced transmission, and increased virulence and pathogenicity, including in mammals^23,24,27^. However, we did not find mutations in PB2 (E627K, D701N, K702R) that have been specifically linked to mammalian host adaptation and enhanced transmission^24,26^. In fact, the viruses sequenced from sea lion and dolphin were genetically similar to each other, but also to viruses from all positive birds, underscoring that it is not possible to discern the direction of transmission among these species from the data available.

However, our phylogenetic analysis does support a single introduction of 2.3.4.4b into Peru from North America, presumably through the movements of migratory wild birds that travel south during the boreal winter, setting the stage for infection of local sea birds that share habitats with marine mammals.

There are multiple possible transmission routes for local transmission among species that involve direct contact or indirect environmental transmission. For one, seabirds share feeding spaces with both sea lions and dolphins, providing ample opportunities for direct contact between animals at sea ^28-31^. Direct contact also occurs on islands, islets, and guano headlands, especially in protected areas where large and dense breeding colonies of sea lions and seabirds cohabitate, and where indirect transmission is also possible via guano runoff into the surrounding waters^28,29,31,32^. Another scenario for transmission involves carnivory and scavenging of infected animal carcasses by marine and terrestrial carnivores, as well as by raptors, gulls, and other scavenger birds^2,33,34^. Fishing docks, where fishermen often dispose of waste by dumping it at sea, attract seabirds, sea lions, marine otters and others that come to feed. Many docks along the Peruvian coast also function as tourist attractions, where seabirds and sea lions are purposely fed to create photo opportunities, building large congregations of wild animals that also increase the risk of transmission to humans. The confirmed presence of HPAI A/H5N1 in 2 species of resident guano seabirds, the Peruvian pelican and Guanay cormorant, provides another potential route for future transmission to humans^35,36^, as guano is widely used to fertilize crops. Finally, the Peruvian dessert coast is home to large poultry operations that hold millions of chickens, which makes them especially susceptible to viruses that circulate in adjacent wild and migratory birds.

There are outstanding questions about which migratory bird species are involved in the long-distance dissemination of HPAI from North to South America, possibly by way of Central America. We detected clade 2.3.4.4b in a migratory sanderling (*Calidris alba*) that would have arrived in Peru after breeding in the Canadian arctic. However, *Calidris* spp. are an unlikely conduit for HPAIV A/H5N1 because experimental inoculations result in death or disease within 5 to 11 days of inoculation^37,38^.

Given the unlikelihood of a successful long-distance migration for a clinically infected bird, we suspect the sanderling was infected locally. Our phylogenetic analysis supports multiple independent introductions of HPAI from North America into South American countries for which sequence data was available at the time of this study, including Peru, Ecuador, and Venezuela. This contrasts with the single introduction of HPAI from Eurasia to North America observed earlier in 2021. Although North America is the primary source of HPAI for South America’s initial HPAI outbreaks, South American countries are likely to become more important sources for each other’s HPAI outbreaks as the virus establishes locally. After observing HPAI A/H5N1 reassort repeatedly within North American viruses, it is possible that the virus will continue to evolve in South America by mutation and reassortment with the genetically distinct South American AIV lineage that is commonly detected in Argentina^39^ and Chile^40^. In addition to the 40 previously characterized variable sites linked to concerning phenotypes that we observed in the 8 Peruvian HPAI A/H5N1 genomes, we identified an additional 30 sites that remain uncharacterized.

There is an urgent need to establish pipelines for efficient real-time genomic sequencing of HPAI to track its evolution as it spreads across Peru and other countries in South America, as well as funding to support characterization of possible new mutations.

The impact of HPAI A/H5N1 on the morbidity and mortality of endangered species like the Andean condor (*Vultur gryphus*)^41^, the Humboldt penguin *(Spheniscus humboldti)*^42^, the marine otter (*Lontra felina*)^43^, and many others is concerning. The Peruvian coast is one of the few places in the world where scavenger condors feed on dead marine animals^44,45^, putting them at risk of infection if they consume contaminated carcasses. The Humboldt penguin lives in large colonies and shares space with guano birds and sea lions, and although the single individual tested here was not positive for HPAI A/H5N1, it is likely a matter of time before we start seeing increased numbers of infected penguins given their proximity to confirmed cases in sea lions and pelicans. Similarly, the marine otter^43^ is an aquatic mustelid that inhabits the same rocky shores of Peru and Chile inhabited by sea lions. HPAI A/H5N1-infected otters have been previously reported^2^, and although none were tested here, we have anecdotal knowledge that some otter carcasses have started to wash onto Peruvian and Chilean shores. Fortunately, marine otters do not live in large groups^46^, which might limit intraspecies contagion.

However, direct mammal to mammal transmission has been suggested as a possible explanation for an outbreak in a Spanish mink farm^6^. The massive South American sea lion die-off currently underway in Peru also supports direct mammal to mammal transmission as a possible viral dissemination route, but confirmation will require further investigations with larger samples sizes and deeper genomic analyses. Peruvian pelicans have also suffered massive die-offs at the beginning of the outbreak in Peru, and now the number of dead Guanay cormorants and Peruvian boobies is starting to increase as well. Peruvian pelicans are considered near-threatened worldwide^47^, but in Peru they are classified as endangered due to recent large population declines resulting from severe El Niño events associated to global warming and to overfishing of their main source of food, the Peruvian anchovy^12^. For these reasons, efforts should be made to assess the impact of ongoing mass mortalities on Peruvian marine sea birds and mammals.

Finally, an even larger concern is the possibility of spillover into human populations, as has been already documented^48,49^, followed by massive human-to-human transmission. Previous human cases have resulted in fatalities^50^, which has led the WHO to declare that the current zoonotic threat from HPAI A/H5N1 remains elevated and that member states should remain vigilant and consider mitigation steps to reduce human exposure^51^. In Peru, the current outbreak is occurring along the Pacific coast and during the austral summer, when many people go to the beach. It is not uncommon for uninformed beachgoers (and their pets) to interact with sick and disoriented animals without any knowledge of the risks and without personal protective equipment, or for free-roaming dogs in rural and semi-rural coastal areas to encounter sick or dead animals as they scavenge for food. This has led government authorities to relocate live animals that show up in places where they do not belong, or euthanize sick individuals and appropriately dispose of their carcasses. However, both the pelican and sea lion die-offs have been so massive that it has been very challenging for the authorities to respond in a timely manner. More public awareness campaigns are needed, and they should focus on educating the public to avoid contact with infected animals^52^. Animal workers, particularly municipal personnel tasked with cleaning duties, need additional training in the proper use of personal protective equipment, and on management and disposal of infected carcasses^13,52^. Syndromic surveillance of personnel dealing with deceased animals, coupled with PCR-based diagnostics and/or serology of suspected cases, is also highly recommended for early detection of clinical infections and transmission. These are all areas of opportunity to strengthen ties between governmental institutions at all levels, expert researchers within academia, and the larger scientific community operationalizing a One Health approach. The work reported here exemplifies such collaboration, and although many of the variable sites identified in Peruvian H5N1 viruses will need further validation with both downstream *in-vitro* and in-*vivo studies*, the fact that this is the first report of HPAI A/H5N1 in marine birds and mammals in Peru represents a significant contribution to the study of a highly pathogenic virus with serious pandemic potential for humans.

## Acknowledgments

The authors thank Sherilym Castillo, Esthefany Ruíz, Pedro Ramírez, Max Guerra, Antero Martínez, Wendy Rojas, Pedro Ramírez, Max Guerra, Antero Martínez, Renato Colán, José Cerón, and Karina Espinoza from SERFOR for support during field work; Julio Reyes and Joe Macalupú from IMARPE for support with dolphin species identification and sampling permissions, respectively; Daniel Alama and Paul Cieza from SENASA for support with sample conservation and transport from field to lab; Miryam Quevedo from UNMSM for providing equipment for fieldwork; and Rosa Vento and Jorge Martínez from WCS for general logistical support.

## Ethical Statement

Sample collection was carried out by employees of the Servicio Nacional Forestal y de Fauna Silvestre (SERFOR) of the Peruvian Ministerio de Desarrollo Agrario y Riego (MIDAGRI) as part of their official duties under permit # XX, along with veterinarians from the Wildlife Conservation Society Peru (WCS).

## Funding Information

Authors are employees of the Peruvian Ministerio de Desarrollo Agrario y Riego (MIDAGRI), of the Pontificia Universidad Catolica del Peru (PUCP), of the Wildlife Conservation Society, of the University of California – Davis, and of the US Government. This work was prepared as part of their official duties. M.L. has additional support from the National Institute of Allergy and Infectious Diseases at the National Institutes of Health under award U01AI151814. This work was also supported by the Centers of Excellence for Influenza Research and Response, National Institute of Allergy and Infectious Diseases, National Institutes of Health (NIH), Department of Health and Human Services, under contract 75N93021C00014, and by the Intramural Research Program of the US National Library of Medicine at the NIH. The funders had no role in study design, sample collection, data collection and analysis, decision to publish, or preparation of the article.

**Supplement Table 1:**
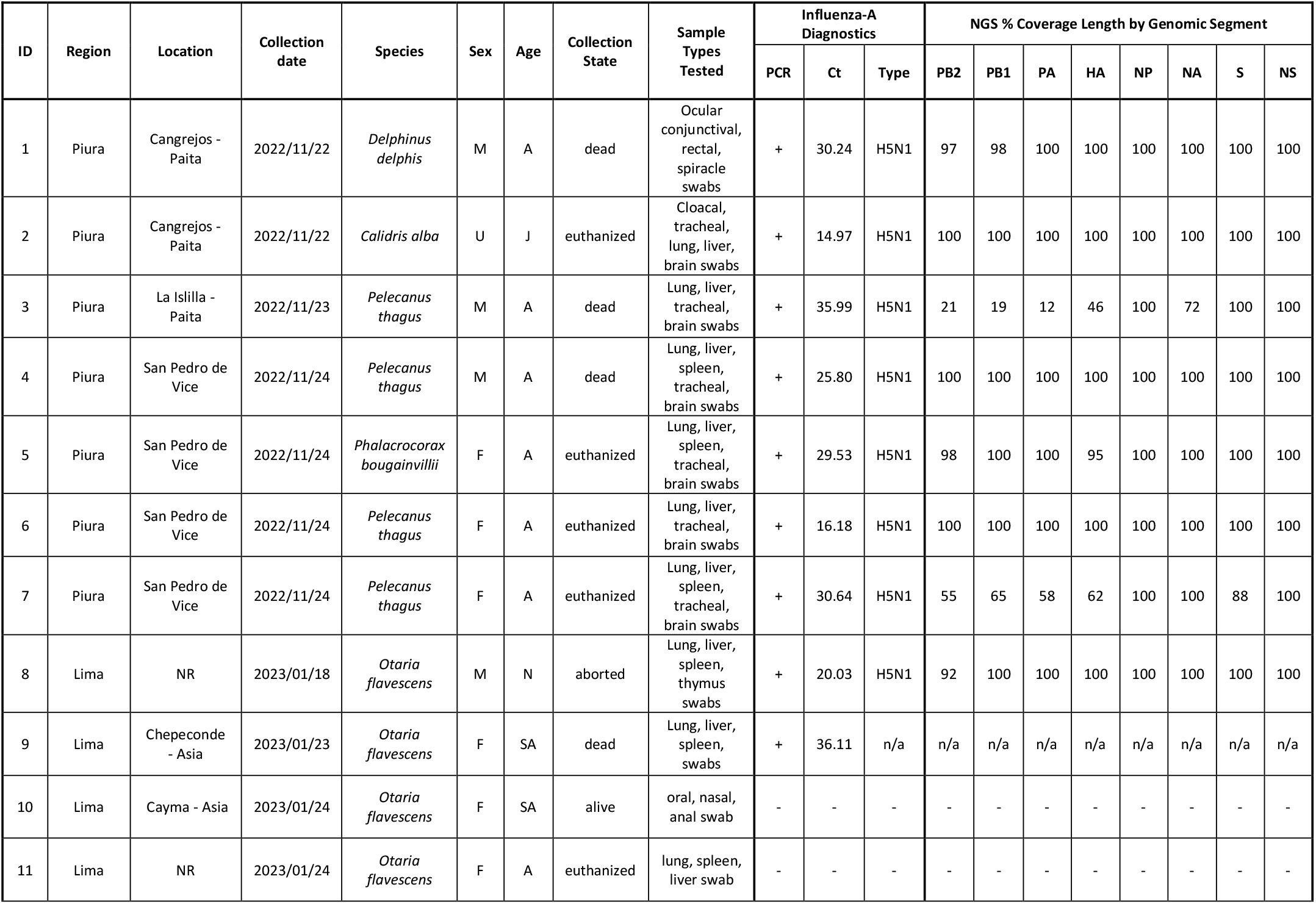
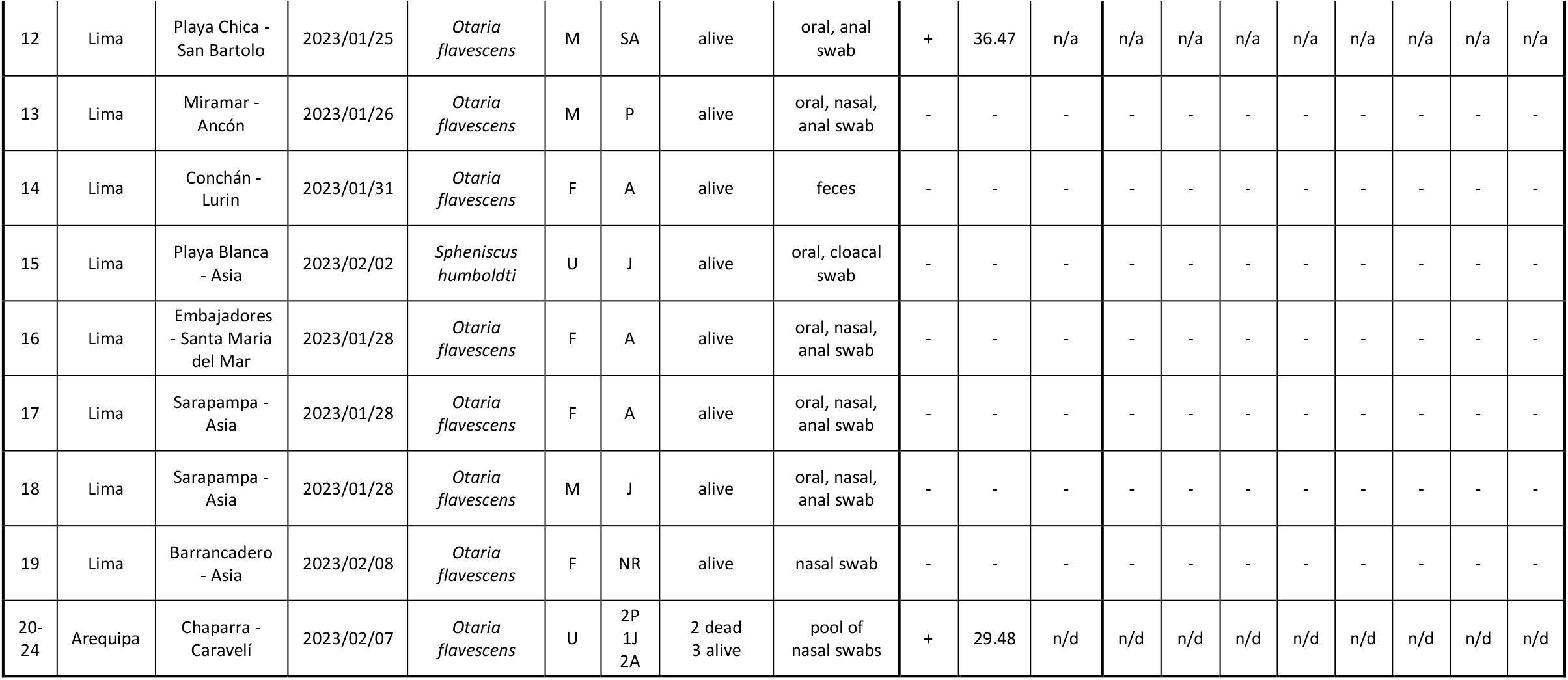
Sampled animals and testing results, including genome coverage information by segment. NR: not recorded, M: male, F: female, U: unknown, A: adult, SA: sub-adult, J: juvenile, P: pup, N: newborn, n/a: not applicable, n/d: not done.

**Supplement Table 2:**
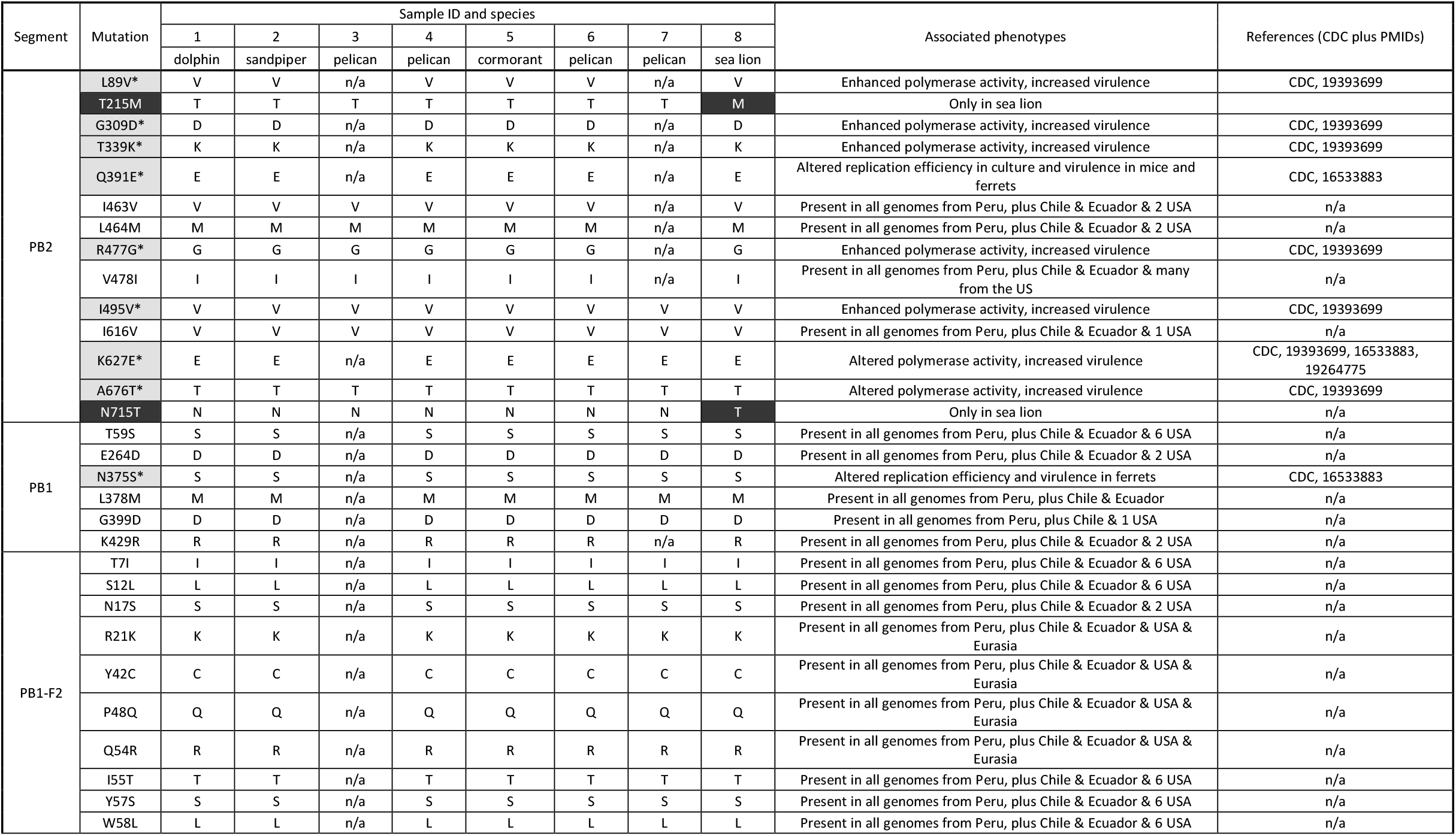
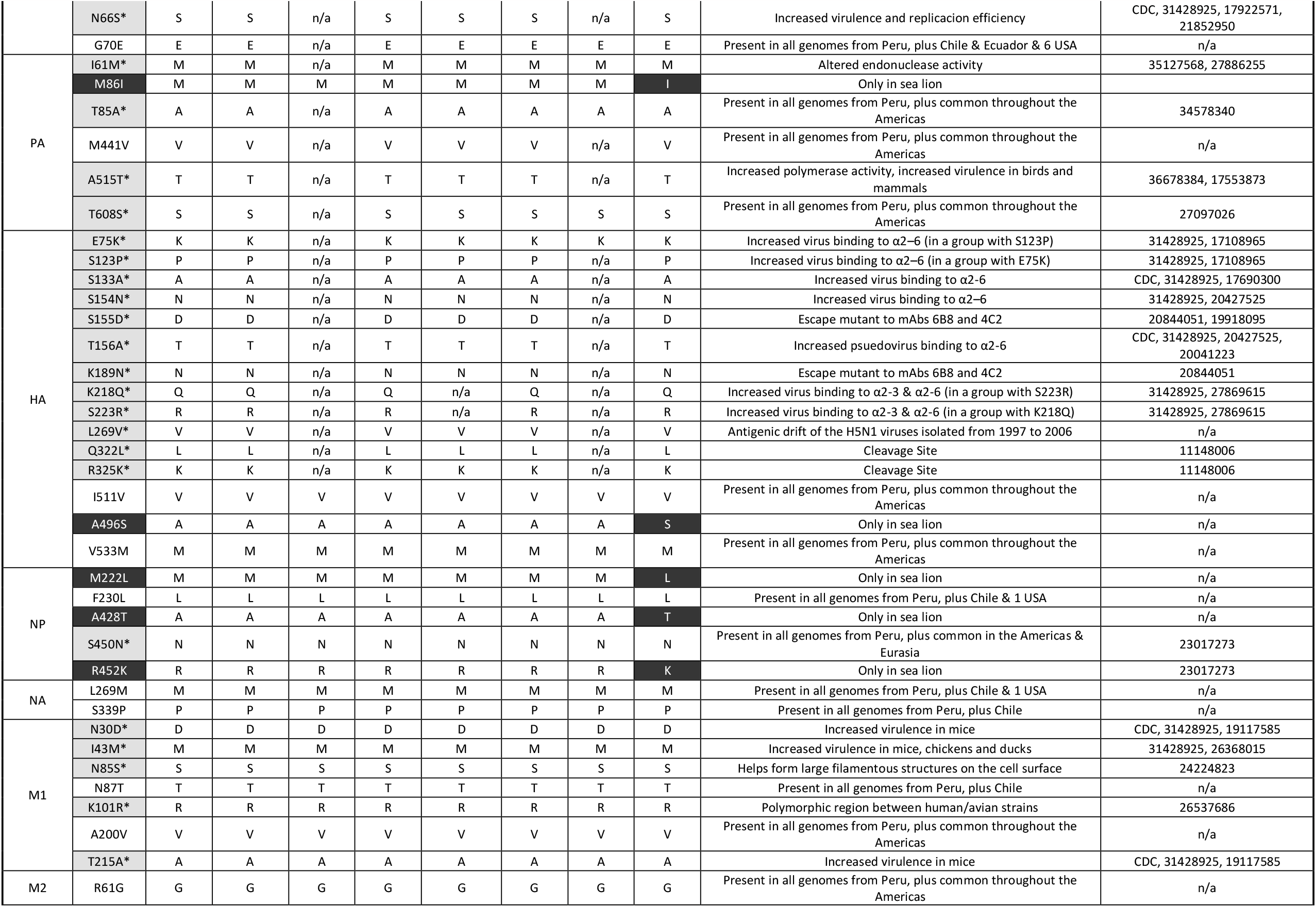
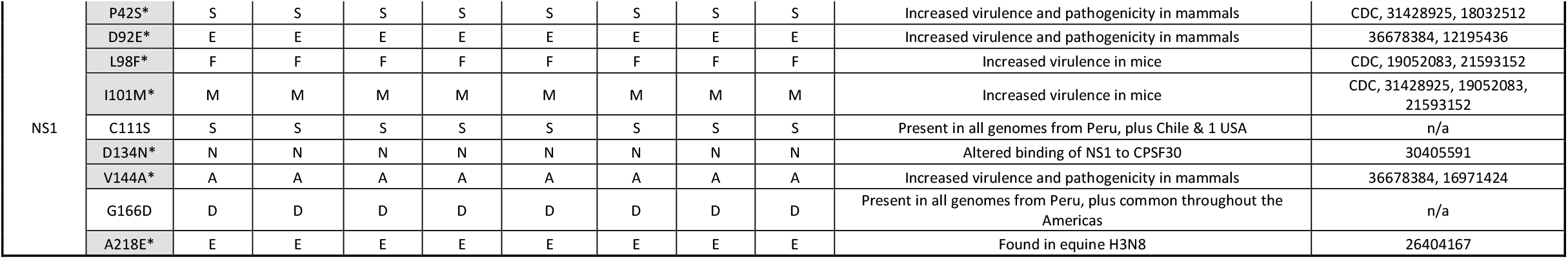
SNP and mutational analysis of Peruvian HPAI a/H5N1 viruses. More than 70 variable sites were identified relative to the original A/H5N1 goose/Guangdong reference from 1996 and the A/Vietnam/1203/2004 reference used to annotate amino acid positions in the CDC inventory^23^. 40 mutations identified have been previously reported as associated with mutations of interest and are shown shaded in gray with an asterisk (*). The remaining 30 mutations have not been previously characterized. An additional 7 mutations only present in sea lion (shaded black) that may be of interest provided they are confirmed in other mammals.

**Supplement Figure 1:**
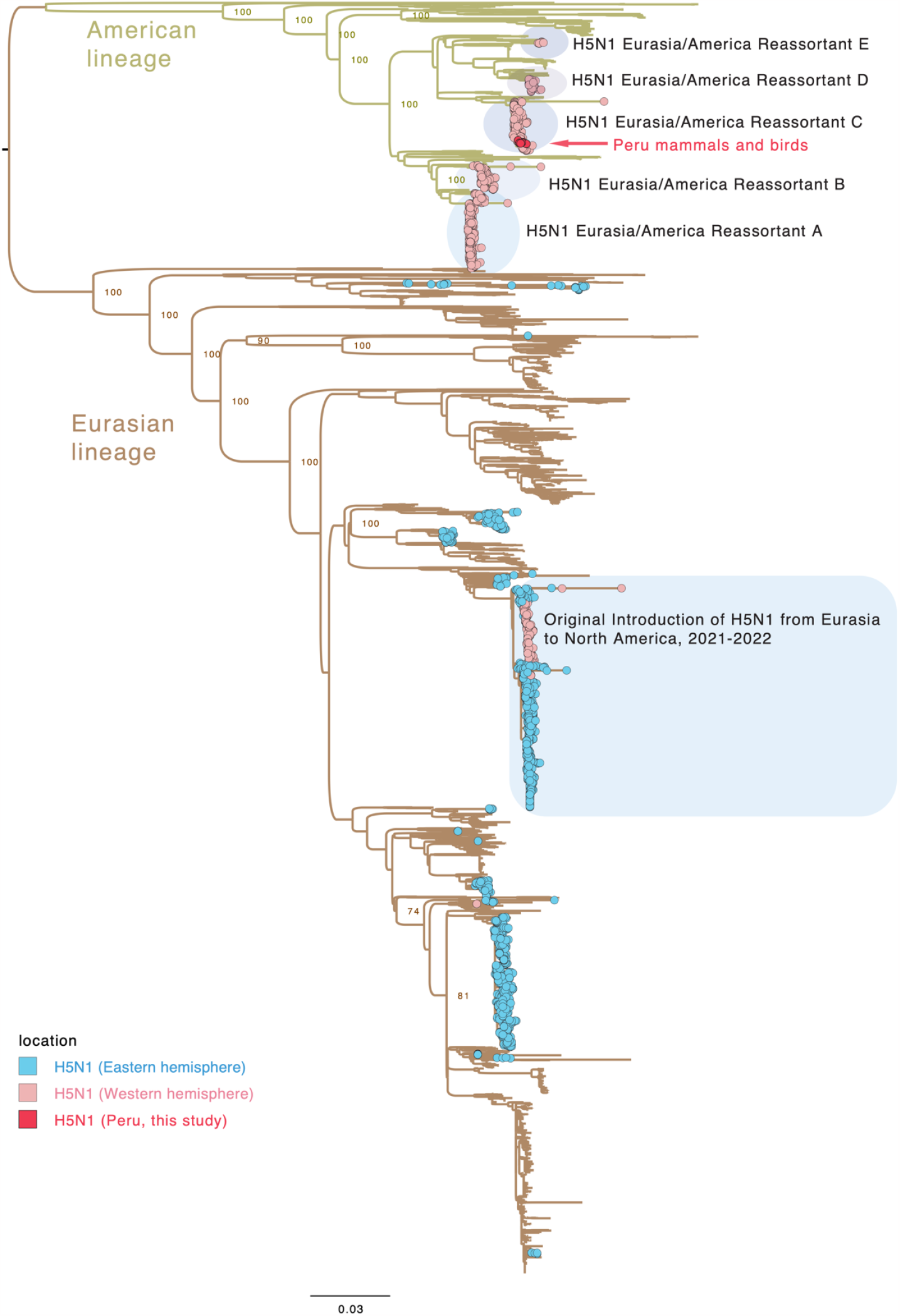
Global H5N1 phylogeny based on the PB2 segment.

**Supplement Figure 2:**
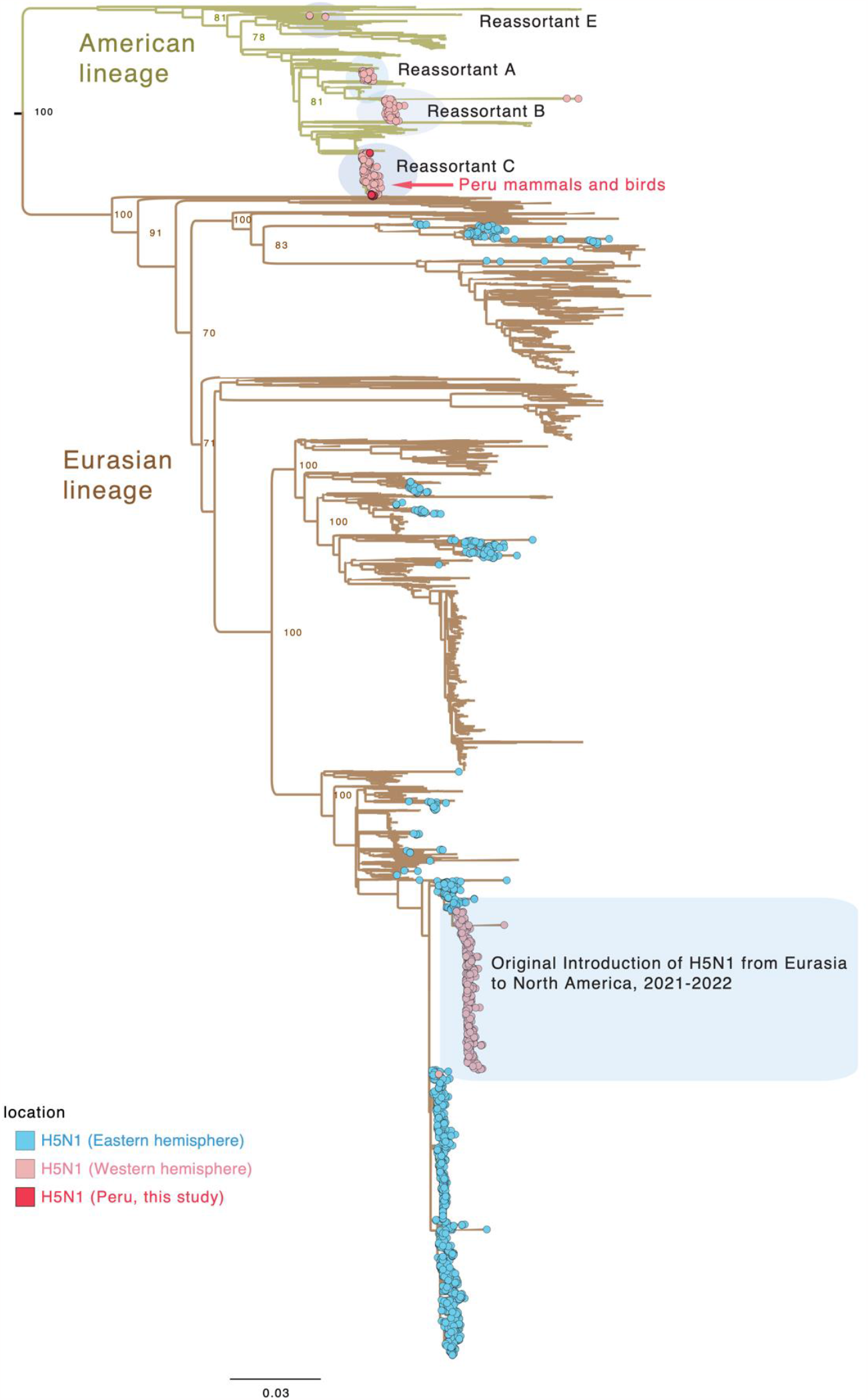
Global H5N1 phylogeny based on the PB1 segment.

**Supplement Figure 3:**
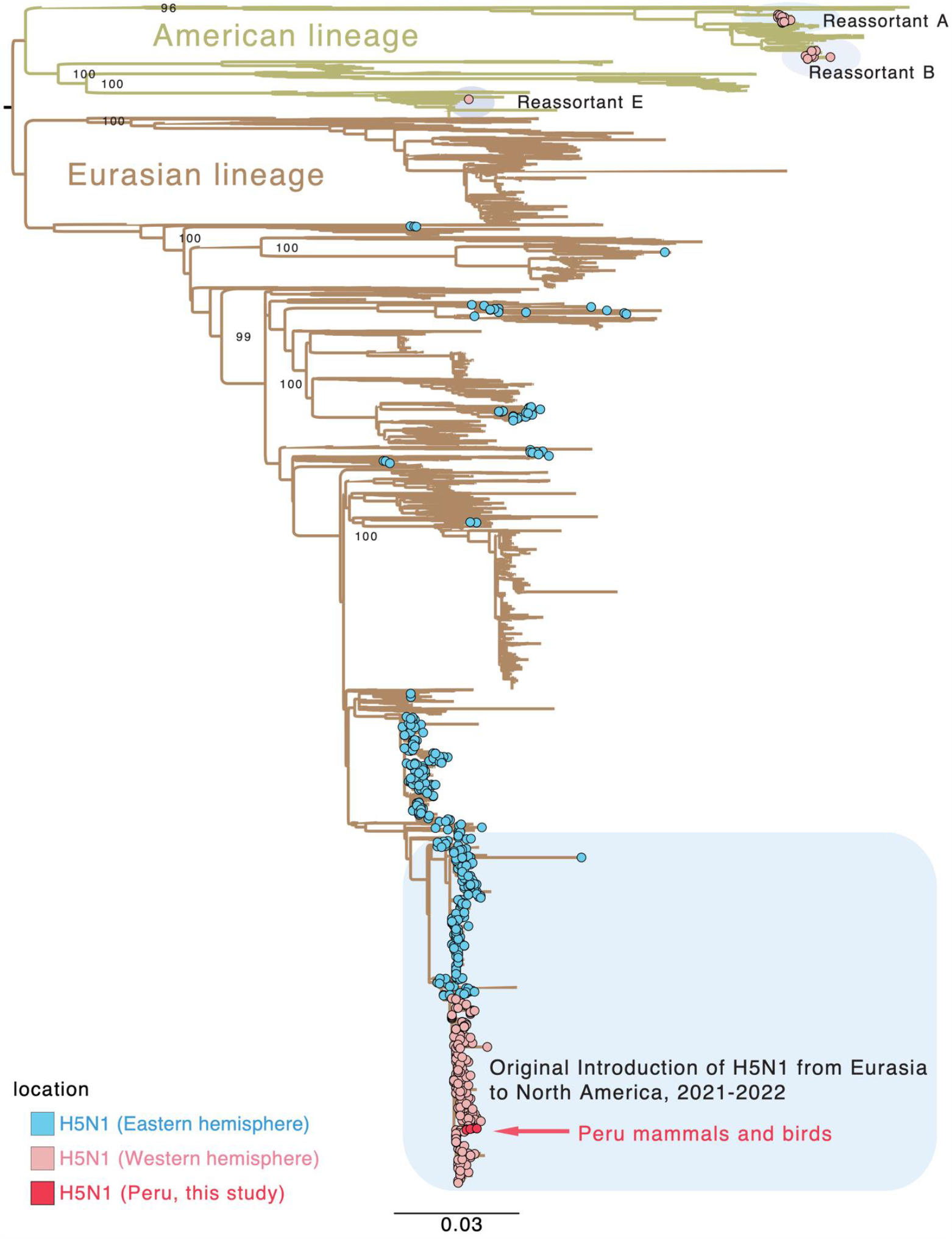
Global H5N1 phylogeny based on the PA segment.

**Supplement Figure 4:**
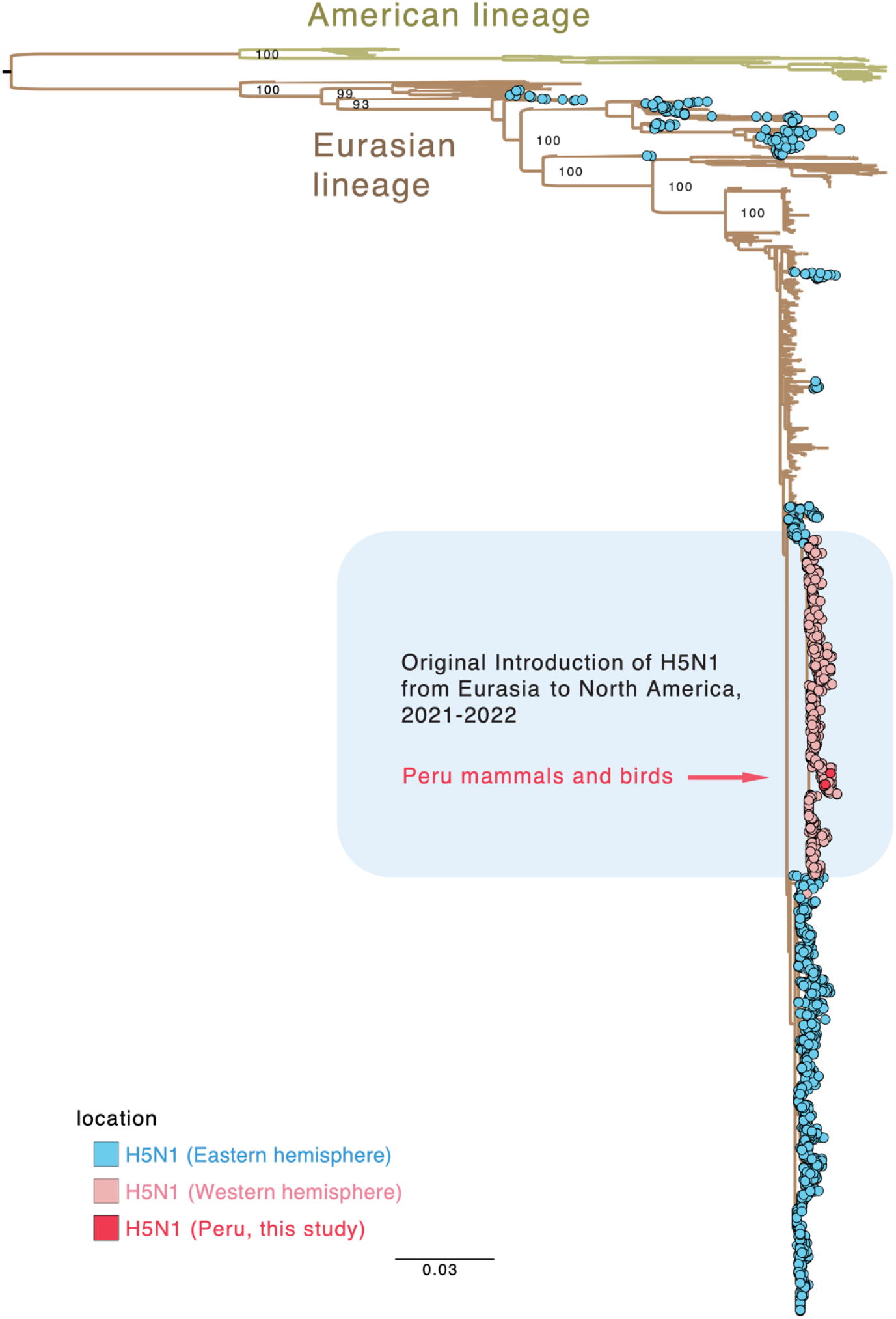
Global H5N1 phylogeny based on the HA segment.

**Supplement Figure 5:**
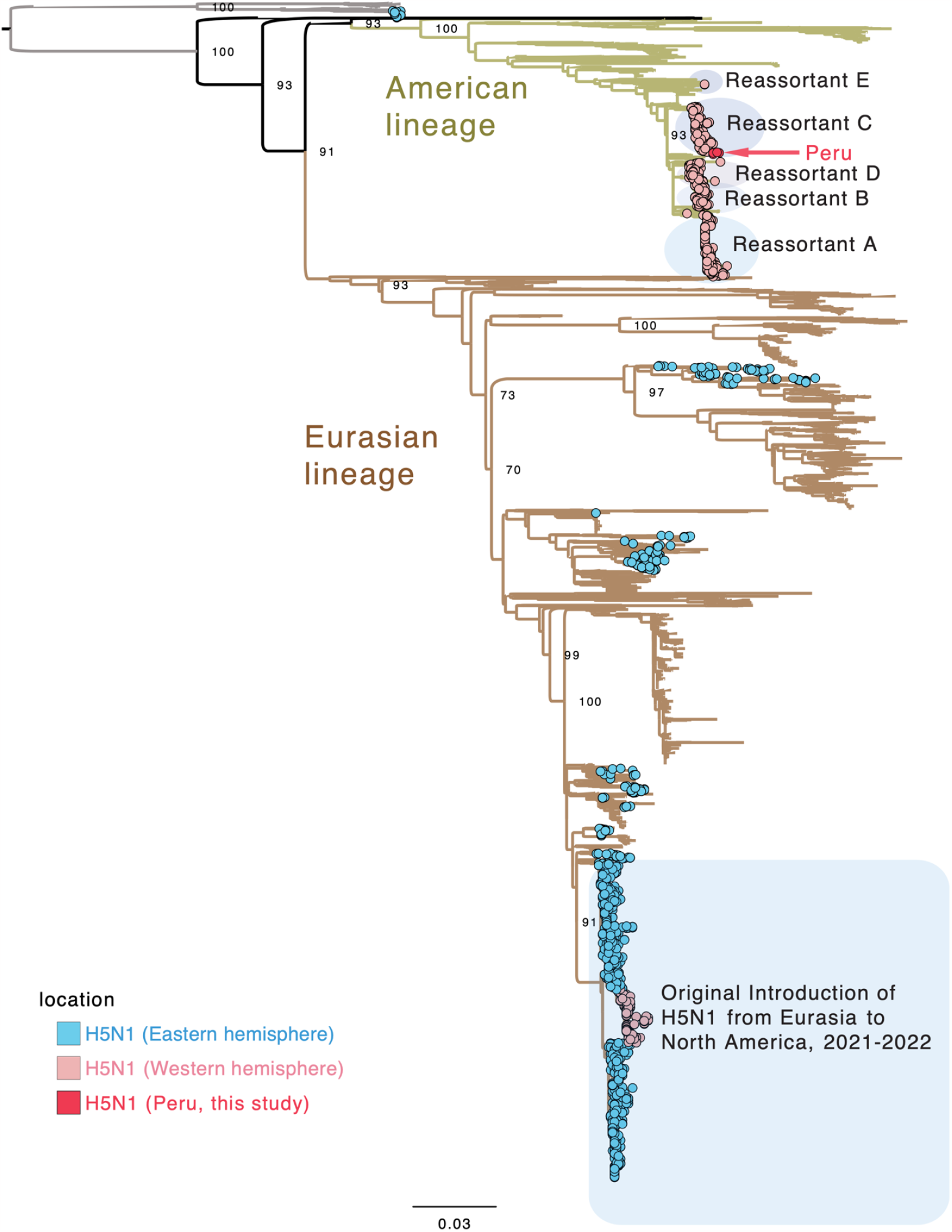
Global H5N1 phylogeny based on the NP segment.

**Supplement Figure 6:**
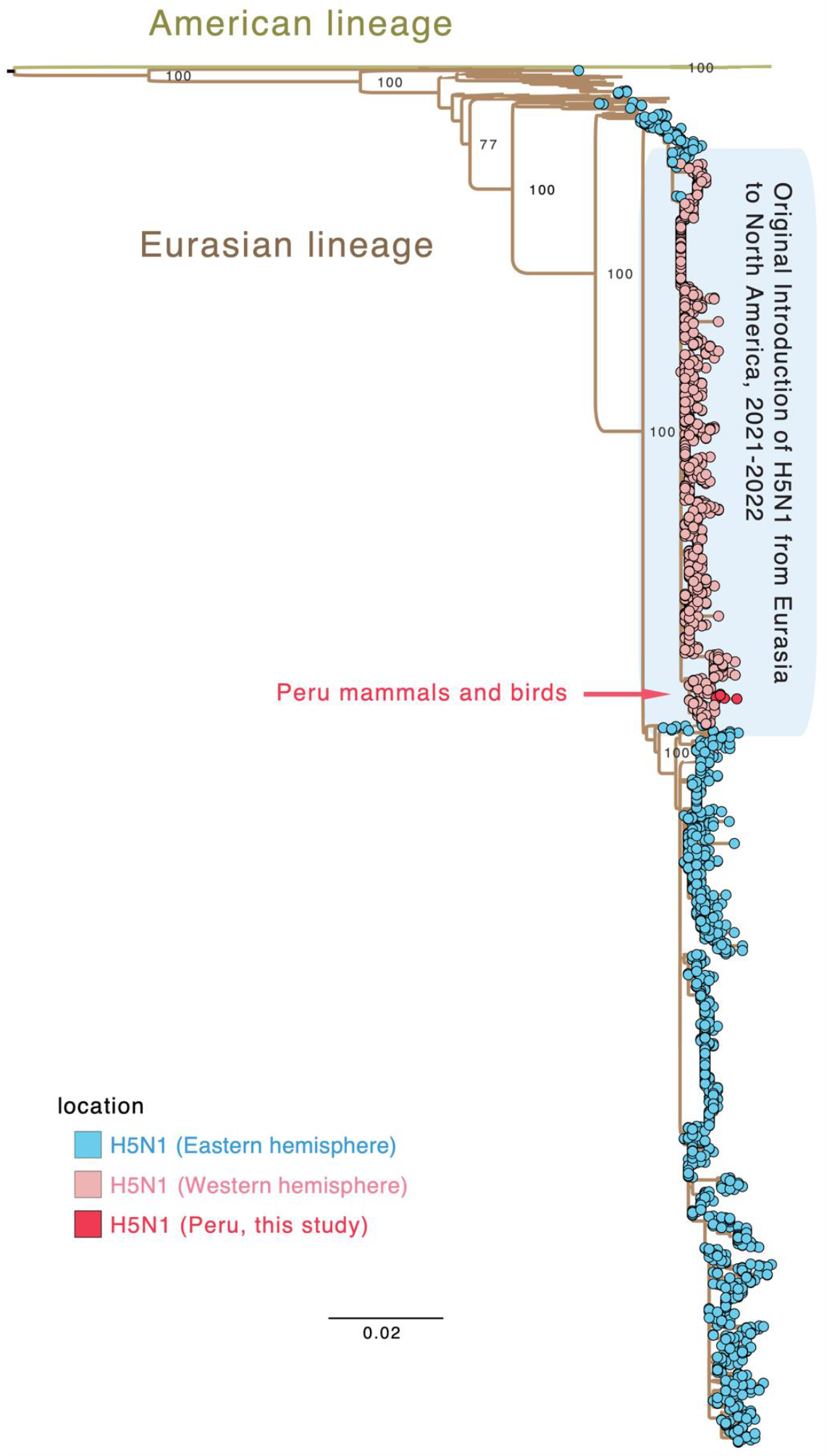
Global H5N1 phylogeny based on the NA segment.

**Supplement Figure 7:**
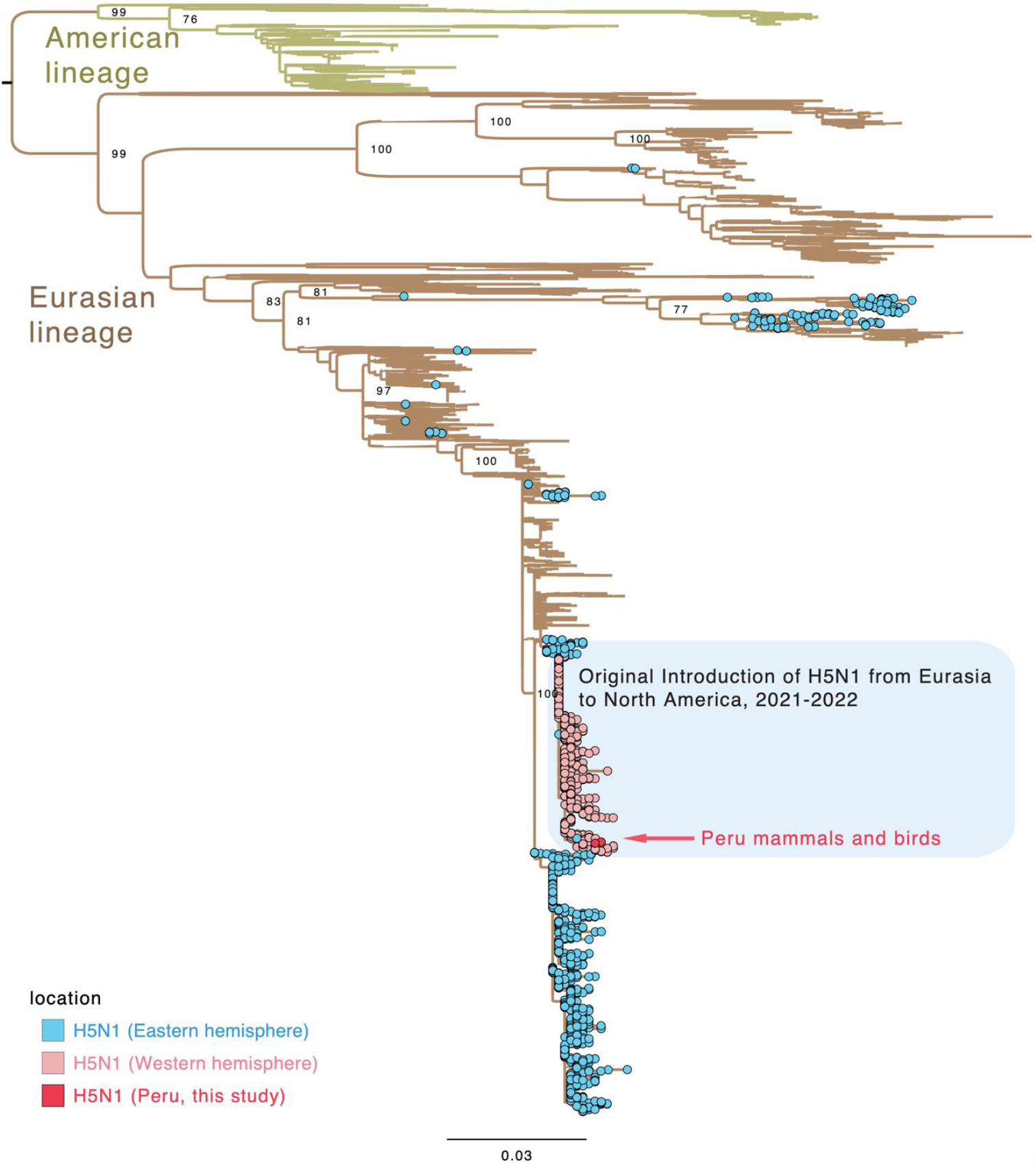
Global H5N1 phylogeny based on the MP segment.

**Supplement Figure 8:**
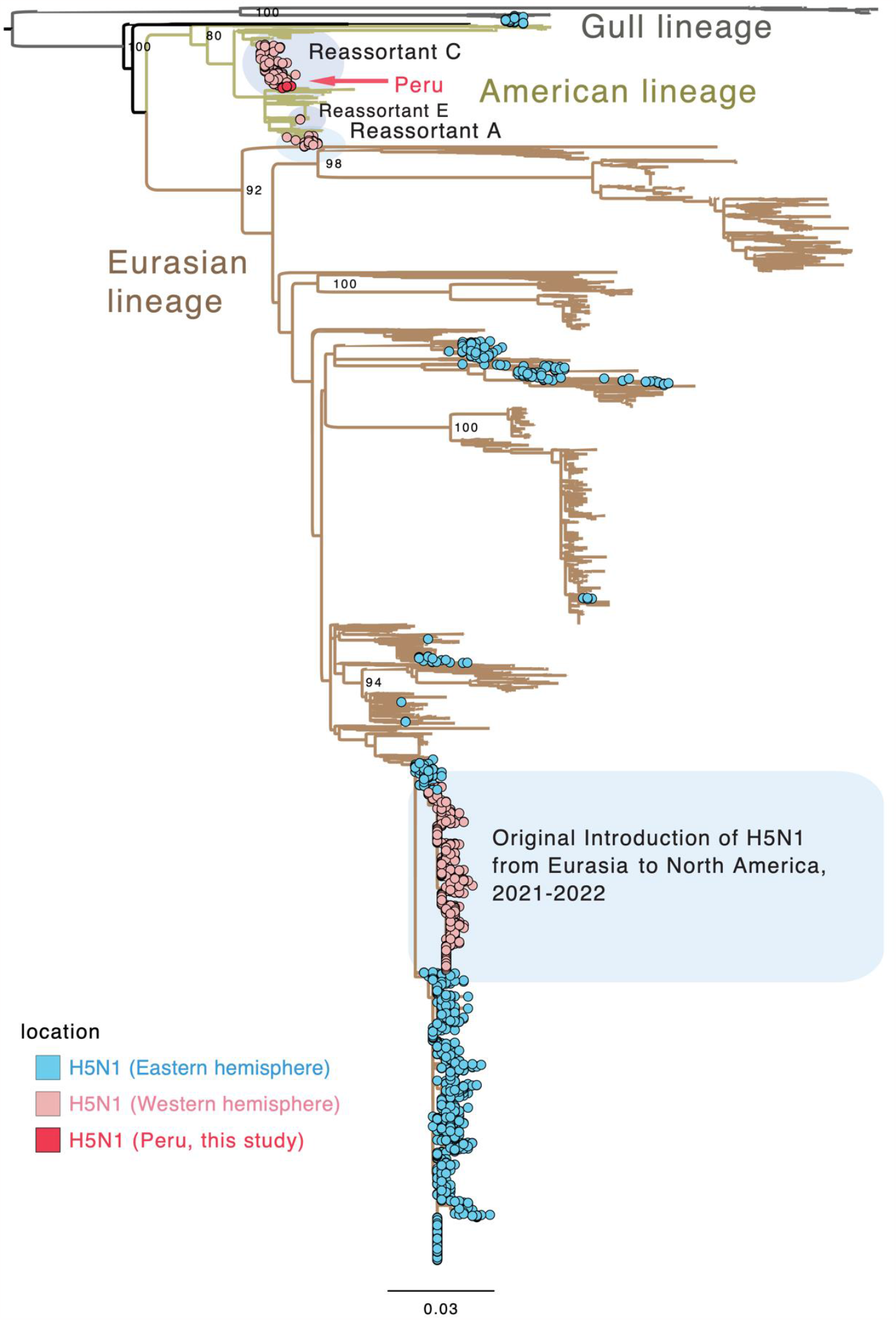
Global H5N1 phylogeny based on the NS1 segment.

